# Quality Matters: Deep Learning-Based Analysis of Protein-Ligand Interactions with Focus on Avoiding Bias

**DOI:** 10.1101/2023.11.13.566916

**Authors:** Manuel S. Sellner, Markus A. Lill, Martin Smieško

## Abstract

The efficient and accurate prediction of protein-ligand binding affinities is an extremely appealing yet still unresolved goal in computational pharmacy. In recent years, many scientists have taken advantage of the remarkable progress of deep learning and applied it to address this issue. Despite all the advances in this field, there is increasing evidence that the typically applied validation of these methods is not suitable for medicinal chemistry applications. This work assesses the importance of dataset quality and proper dataset splitting techniques demonstrated on the example of the PDBbind dataset. We also introduce a new tool for the analysis of protein-ligand complexes, called po-sco. Po-sco allows the extraction of interaction information with much higher detail and comprehensibility than the tools available to date. We trained a transformer-based deep learning model to generate protein-ligand interaction fingerprints that can be utilized for downstream predictions, such as binding affinity. When using po-sco, this model generated predictions that were superior to those based on commonly used PLIP and ProLIF tools. We also demonstrate that the quality of the dataset is more important than the number of data points and that suboptimal dataset splitting can lead to a significant overestimation of model performance.

## 1 Introduction

Protein-ligand interactions, which drive molecular recognition processes play a crucial role in many biological processes. The accurate prediction of binding affinities associated with such interactions belongs to the most important tasks in drug design. In recent years, various computational methods have been conceived and developed to predict protein-ligand binding affinities, which can help guide drug discovery efforts and reduce the time and cost associated with experimental screening [1–11].

### 1.1 Prediction of binding affinities

A well-established approach to create protein-ligand binding modes is molecular docking. This method employs algorithms that search for the optimal orientation and conformation of the ligand (referred to as pose) within the active site of the protein and then try to quantify the binding energy based on the interaction between the protein and the ligand [12–16]. While it is relatively simple and fast, it is not very accurate in predicting binding affinities [17–20]. Another method that has gained increasing importance during the last decade is free energy perturbation (FEP). It relies on molecular dynamics simulation to calculate the change in free energy upon binding of the ligand to the protein. This method can deliver predictions of relative and absolute binding affinities of chemical accuracy, but is computationally extremely demanding [6, 21–24]. Other computational methods for predicting binding affinities include molecular mechanics Poisson-Boltzmann surface area (MM-PBSA) and molecular mechanics generalized Born surface area (MM-GBSA), which involve the calculation of binding free energies using molecular mechanics force fields and implicit solvation models [25–27]. The development of these computational methods has facilitated the prediction of protein-ligand binding affinities and has the potential to accelerate drug discovery and design efforts. However, most of these methods still suffer from either poor accuracy or high computational cost.

In recent years, deep learning approaches have gained popularity for predicting protein-ligand binding affinities. Deep learning models use artificial neural networks to learn the complex relationships between the molecular features of the protein and ligand and their binding affinity [28]. A commonly used approach is the use of convolutional neural networks (CNNs) to predict binding affinities. These models usually take 1- or 3-dimensional representations of the protein and ligand as input and use convolutional layers to extract local features from the structures. The extracted features are then fed into fully connected layers to predict the binding affinity [29–31]. Although CNNs excel at translational invariance, they are not rotationally invariant, which may cause problems when working with 3-dimensional input data. Another common deep learning approach for the prediction of binding affinities is the use of graph neural networks (GNNs). GNNs operate on molecular graphs, where atoms are considered as nodes and chemical bonds as edges. These models can learn the molecular representation of protein binding sites and ligands from their graph structure. GNNs are commonly used to predict binding affinity by considering binding site residues, ligand atoms, and interactions between them on the molecular graph [32–36]. When implemented correctly, GNNs can have translational, rotational, and reflection equivariance (i.e. E(3) equivariance) [37]. Recently, transformer models [38] have shown great promise in many different fields [39–43]. In protein-ligand interaction prediction, these models take advantage of the power of self- and cross-attention modules to analyze complex interactions between proteins and ligands and predict their binding affinity with seemingly high accuracy [44, 45]. A key advantage of transformer models is their ability to model long-range dependencies in sequence-based data and capture complex interactions between amino acids and ligands. This is critical for accurately predicting binding affinities, as these interactions can be highly specific and context-dependent.

So far, deep learning approaches have shown promising results in predicting protein-ligand binding affinity and have the potential to further advance the field of drug discovery and design. However, the development of accurate deep learning models for predicting binding affinity requires large amounts of high-quality data and careful selection of molecular features to incorporate into the models. In this work, we use a transformer-based model that uses self- and cross-attention to learn from residue-ligand interaction fingerprints to predict binding affinities and protein-ligand interaction fingerprints.

### 1.2 Biases in binding affinity prediction

While deep learning approaches have great promise in predicting protein-ligand binding affinities, they are susceptible to various biases that can limit their accuracy and generalizability.

The data used to develop a deep learning model can be a major source of bias. Such bias can have various origins, for example if the training data used to develop the models are not representative of the true distribution of protein-ligand binding affinity data. This can happen due to, e.g. the limited availability of high-quality binding affinity data, the uneven distribution of binding affinity values, or the use of biased selection criteria for the data. To objectively assess the performance of a model, it is absolutely vital to avoid any overlap between training, validation, and test sets. As a result of such bias, trained models may not be able to accurately generalize to new data outside the training set, leading to overfitting. Fan and Shi have demonstrated that the overlap of proteins and ligands in the test and training set greatly influences the apparent performance of the model and leads to a severe overestimation of the capabilities of the model [46]. They also proposed that such a bias has a greater influence on the performance of a model than its architecture.

Another potential bias in deep learning models is feature bias, where the features used to train the models do not capture all the relevant information about protein-ligand interactions. For example, if the features used only capture the geometric properties of the protein and ligand, they may not account for the dynamic changes that occur during the binding process. Feature bias can also occur if the features used contain too much information, causing the model to overfit. Volkov et al. showed that using 3D information of proteins and ligands often introduces bias if molecular structure information is provided directly to the model [47]. For example, if the fully atomistic structure of the ligand and/or binding site, or the protein sequence is provided to a neural network, it often learns to memorize the specific data and does not generalize well. They proposed that instead of directly using structural information, one should rely only on extracted interactions between the protein and the ligand.

To overcome bias, it is important to carefully curate high-quality datasets that accurately represent the range of binding affinity values, meticulously split the data into training, validation, and test sets, and carefully select the input features of the model.

### 1.3 Types of protein-ligand interactions

The analysis of protein-ligand interactions in a structural complex is critical to understanding the binding mechanism and designing new therapeutics. Thus, it seems logical that a detailed analysis of protein-ligand interactions may aid deep learning models to properly learn binding affinities. One way to analyze these interactions is to identify, enumerate, and quantify specific types of molecular interactions recognized in physics and chemistry, such as hydrogen bonds, salt bridges, hydrophobic interactions, pi-pi, sigma-pi, pi-cation interactions, and halogen bonds.

Hydrogen bonds are arguably the most common and important specific molecular interactions in protein-ligand complexes. These interactions occur between a hydrogen atom carrying a partial positive charge (typically due to an electron-withdrawing carrier atom or functional group) and an electronegative atom featuring a lone electron pair, such as oxygen or nitrogen. The strength of a hydrogen bond depends on the distance and angle between the atoms involved, as well as polarization effects. Analyzing hydrogen bonds can not only provide information on the strength of a ligand binding to a protein, but also on the specificity [48].

Charge-assisted interactions like salt bridges are the strongest type of intermolecular interactions. Salt bridges are formed by the electrostatic attraction of oppositely charged atoms or functional groups. This type of interaction can be crucial to stabilizing the protein-ligand complex if the ligand contains charged functional groups, also due to its long range compared to other interactions [49]. Therefore, recognition of salt bridges, but also potentially unsatisfied charged moieties missing counterparts or even repulsive interactions in a protein-ligand complex is imperative for correctly estimating binding affinities.

Although generally weaker than polar interactions if quantified per interacting atom pair, hydrophobic interactions - due to their numerous occurrence - are a key contributor to the strength of a protein ligand complex [48]. These interactions arise from the close proximity of non-polar atoms that shield each other from unfavorable interactions with water (hydrophobic). Ferreira de Freitas and Schapira found that high-efficiency ligands often show numerous hydrophobic interactions when analyzing protein-ligand interactions in the protein data bank (PDB) [50].

Pi-pi interactions are a specific type of dispersion forces and typically occur between large planar functional groups with pronounced electron delocalization like aromatic rings [50, 51]. These interactions can be important in stabilizing the protein-ligand complex and can provide additional specificity (besides directional H-bonds and salt bridges) for ligand binding [48]. Since pi-pi interactions have been found to be the third most common protein-ligand interaction in the PDB, identifying different forms of pi-pi interactions (such as face-to-face pi-pi and T-shaped sigma-pi) is important for the analysis of binding complexes [50]. Pi-cation interactions are another type of non-covalent interaction that occurs between a positively charged group, such as a metal cation, protonated amine, or guanidinium group, and an electron-rich π-system. Although pi-cation interactions are not as common as pi-pi stacking, these interactions may be important for stabilizing protein-ligand complexes and can also contribute to ligand binding specificity [50].

Halogen bonds are a type of non-covalent interaction that occurs between a halogen atom beyond fluorine (such as chlorine, bromine, or iodine) and either an electrophile or a nucleophile atom depending on the geometry. Most halogen bonds consist of interactions between a region of the halogen with low electron density (called the sigma hole) and electronegative atoms such as oxygen or nitrogen featuring a free electron pair. However, halogen bonds can also form between a halogen atom’s electron-rich belt and electropositive atoms, such as polarized hydrogens. While the former have a linear geometry (i.e. the angle C-X O is close to 180°), the latter halogen bonds prefer a perpendicular setup (i.e. the angle C-X Hpolar is close to 90°) [52, 53]. Thus, as with hydrogen bonds, the strength of halogen bonds depends on the distance and angle between the atoms involved.

In general, analysis of specific molecular interactions in protein-ligand complexes can provide valuable insight into the binding mechanism and can inform the design of novel therapeutics. By understanding the specific interactions involved in protein-ligand binding, researchers can design more effective drugs with higher binding affinity and better selectivity. The detailed analysis of molecular interactions can also help computational scientists develop tools and models for various predictions in the field of drug design or toxicology.

There are a few tools that allow the generation of protein-ligand interaction fingerprints. The two most widely used tools are PLIP [54] and ProLIF [55]. PLIP allows the detection of hydrophobic interactions, hydrogen bonds, aromatic stacking, pi-cation interactions, salt bridges, water-bridged hydrogen bonds, halogen bonds, and metal interactions. The newer ProLIF tool allows for detecting hydrophobic interactions, hydrogen bonds (distinguishing between accepting and donating groups), pi-pi stacking (edge-to-face and face-to-face), pi-cation and cation-pi interactions, salt bridges (with differentiation between cationic and anionic groups in the ligand), donating and accepting halogen bonds, as well as donating and accepting metal interactions. Although these tools are sufficient for the analysis of the most common interactions, they lack more sophisticated functionalities: in particular, the differentiation between backbone and side-chain hydrogen bonds, linear (to nucleophiles) and perpendicular (to electrophiles) halogen bonds, the detection of polarized hydrogen bonds, and repulsive interactions. To address these points, we developed a custom tool for protein-ligand interaction analysis, which is the most sophisticated tool to our knowledge so far. We call our tool po-sco (as an abbreviation of pose-scorer).

### 1.4 The PDBbind dataset

At present, several datasets are commonly used to train artificial intelligence models to predict protein-ligand binding affinities. These datasets vary in size, diversity, and quality, and each has its own advantages and disadvantages.

One of the most widely used datasets is the PDBbind database, which harbors experimentally determined binding affinities for a diverse set of protein-ligand complexes [56]. The PDBbind database has been used to train and evaluate a wide range of machine learning and deep learning models, and is frequently used as a benchmark to assess the performance of new models [2, 57–60]. One advantage of the PDBbind database is its size and diversity, which allows for the development of models that can potentially generalize well to new protein-ligand complexes. Another strong feature of the PDBbind dataset is that it links the three-dimensional protein-ligand structural information (including their PDB ID) with the corresponding binding affinity data. This allows the use of three-dimensional features, such as molecular interactions, in prediction models.

The major disadvantage of the PDBbind database is that it contains a relatively small number of high-quality complexes. This may be limiting for models that need a high level of detail. Also, it is not trivial to achieve a good dataset split limiting the bias as much as possible. The PDBbind dataset is divided into a general, refined, and core set. The general set represents the largest part and contains complexes of lower quality. The refined set is a curated set that contains only complexes that meet certain quality criteria. These criteria include quality checks of the 3D structures (e.g., a resolution *<* 2.5 Å, no covalently bound ligands, and no steric clashes) and the binding data (e.g., K_d_ and K_i_ instead of IC50, exact values instead of ranges, and affinity data only within a pharmaceutically relevant range). The core set is the smallest of the three and contains only high-quality complexes of high diversity. Some researchers use the general, refined, and core set as training, validation, and test set, respectively [2]. Others randomly split one of the sets to construct the datasets used for training [59]. Since there are many overlaps of proteins (i.e. Uniprot IDs) and ligands (with the same 2D structure) among the different sets of PDBbind, both approaches probably lead to the introduction of a large bias, as discussed in Section 1.2. Thus, a great deal of attention must be paid to using the PDBbind dataset for developing tools for protein-ligand binding affinity prediction.

The PDBbind dataset contains a multitude of protein-ligand complexes. Each complex is provided in the form of a protein structure file, a file containing only the binding pocket of the protein, and the ligand in two different formats. While the ligand file contains all explicit hydrogen atoms, the protein structure contains only polar hydrogen atoms, and water molecules contain no hydrogen atoms. This may be problematic, e.g., because the orientation of co-crystallized water molecules is unknown, although water molecules are considered imperative for protein-ligand binding [31, 61–63]. In addition, the ligand structures in the PDBbind dataset are provided in neutral state. Thus, the generation of the correct protonation states of the ligand (and the protein) at the desired pH must be performed by the user.

### 1.5 Our contribution

In this work, we developed an in silico tool called po-sco to comprehensively analyze native protein-ligand interactions as they appear in crystals and create residue-based interaction fingerprints. These fingerprints are designed to be constructed in a way that excludes any explicit information about the ligand structure or binding site geometry in order to avoid any structural bias. An attention-based deep neural network is then employed to learn the corresponding experimentally determined protein-ligand binding affinities. Furthermore, the model converts residue-based interaction fingerprints into a single protein-ligand interaction fingerprint that can be used for downstream tasks.

As we use the PDBbind dataset to train and validate our model, we also show how different splitting procedures affect the apparent model performance, while already having reduced structural bias by focusing on molecular interactions rather than fully atomistic structures.

With this work, we hope to shed light into the murky waters of deep learning-based protein-ligand binding affinity prediction and highlight the importance of high-quality datasets and features.

## 2 Results and Discussion

### 2.1 Po-sco interaction analysis

Po-sco provides a great level of detail when it comes to the analysis of protein-ligand complexes. It is capable of detecting the large majority of currently recognized types of intermolecular interactions found in protein-ligand complexes. Additionally, its detection algorithms implement a high degree of physics and medicinal chemistry rules to provide the user with very fine-grained analyses. While traditional protein-ligand interaction analysis tools are limited to binary fingerprints, po-sco extends binary fingerprints with continuous values, allowing even more information to be stored in a single fingerprint. The combination of all these features leads to 28 binary and 6 continuous features extracted by po-sco (see Table 1 for a full list of included features).

**Table 1:**
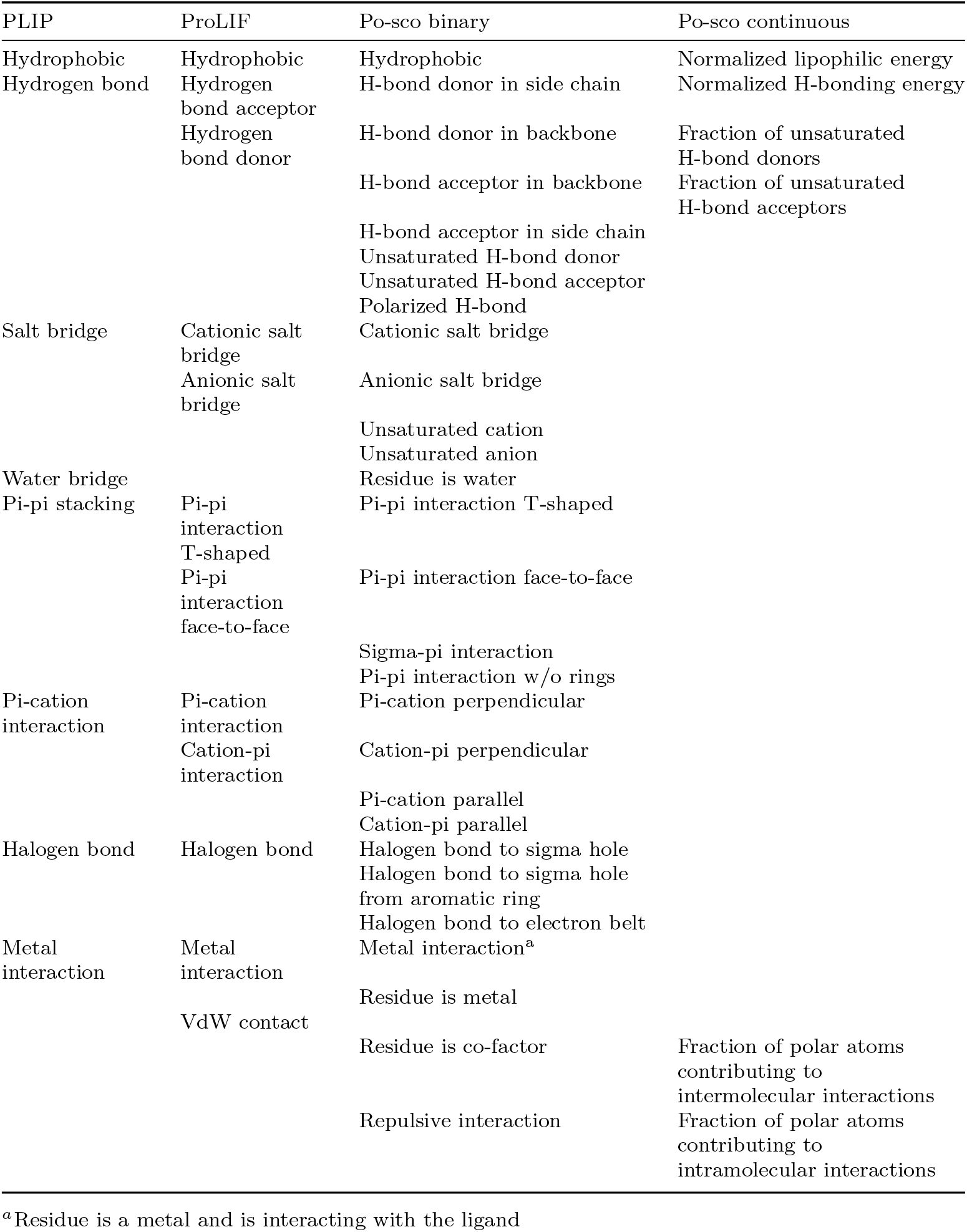
Protein-ligand interactions calculated by PLIP, ProLIF, and po-sco. Where not otherwise stated, the interactions are listed from the perspective of the protein residue. Except for the interaction types under “Po-sco continuous”, the presence or absence of an interaction is indicated by a binary value. For “Po-sco continuous”, the values are continuous, ranging from 0 to 1.

To demonstrate the binary interactions identified by po-sco, we analyzed the crystal structure of human tryoptophan hydroxylase type 1 co-crystallized with the ligand LP-533401 (PDB ID 3HF8). An overview of the identified interactions can be found in Figure 1. For better visibility, we split the interactions in polar (Figure 1A), hydrophobic (Figure 1B), and exotic (Figure 1D) interactions. Furthermore, we show the identified unsaturated polar functional groups in Figure 1C. The figure shows that po-sco was able to reliably identify the most important interactions in high detail. We believe that not only the detailed analysis of the existing interactions, but also the information of missing interactions as shown in Figure 1C are important for assessing the quality of a protein-ligand complex.

**Fig. 1:**
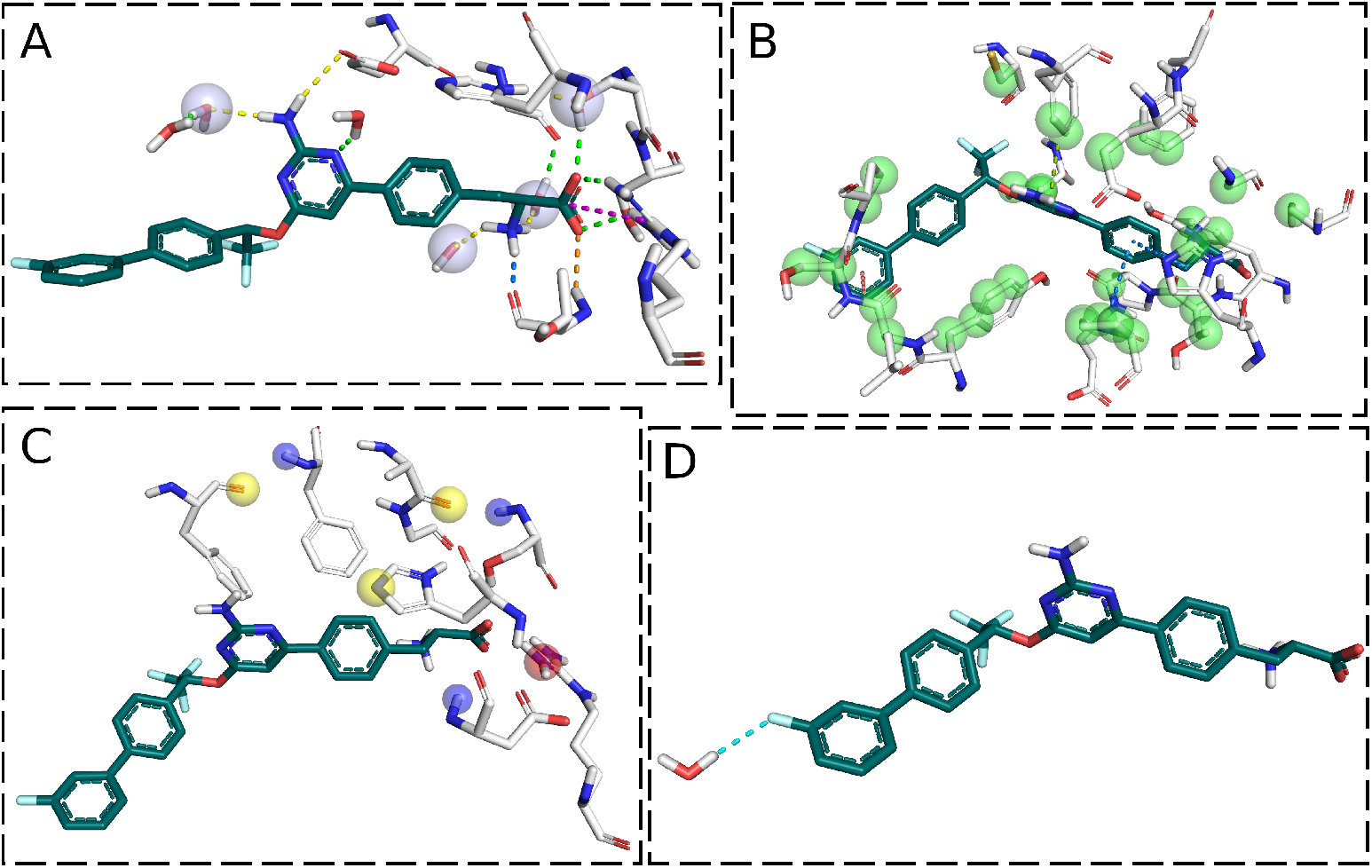
Visualization of the binary interactions identified by po-sco in PDB ID 3HF8. A) Polar interactions where yellow, green, blue, and orange dashed lines represent interactions with H-bond acceptors in side chains, H-bond donors in side chains, H-bond acceptors in the backbone, and H-bond donors in the backbone, respectively. Salt bridges are shown as dashed magenta lines, and polarized H-bond acceptors are marked with a semi-transparent sphere. B) Hydrophobic interactions where blue, green, and orange dashed lines represent face-to-face pi-pi, T-shaped pi-pi, and pi-pi interactions to smaller elements, respectively. Green spheres mark residues with hydrophobic contacts with the ligand. C) Unsaturated polar functional groups. The yellow, blue, and red spheres represent unsaturated uncharged acceptors, unsaturated uncharged donors, and unsaturated cationic groups, respectively. D) A single exotic group (halogen bond to electron-rich belt) is shown.

### 2.2 Attention-based prediction model

The first goal of this work was to find out if more detailed interaction analysis benefits the prediction of binding affinities. As shown in Table 1, the level of detail provided by PLIP, ProLIF, and po-sco differs significantly. Therefore, we trained multiple versions of the same model with interaction fingerprint data extracted from the three analytic tools. For po-sco, we trained two models, one with explicit information on the ligand happiness and one without it. In order to compare the model performance, we calculated the Pearson correlation coefficient (PCC), the mean unsigned error (MUE), and the root mean squared error (RMSE) between predicted and true, experimentally determined affinities. It is important to note here that the goal was not to get a perfect prediction of binding affinities, but to compare how different model input data affect the quality (predictivity) of results while using the same model architecture.

Table 2 serves as the performance comparison of the different models. All three performance metrics are worst for PLIP and best for po-sco (with ligand information). They also clearly correlate with the level of detail provided by the different tools. ProLIF distinguishes between hydrogen bond donors and acceptors as well as between anionic and cationic salt bridges. These details are missing in PLIP. Although ProLIF lacks detection of water bridges, it performed better than PLIP. This may be due to the lower occurrence of these kinds of interaction and, therefore, to their lower relative importance for the prediction of binding affinities. Hydrogen bonds and salt bridges, on the other hand, are among the most common protein-ligand interactions and are therefore considered highly important. Thus, it seems reasonable that a higher level of detail in these common interaction types leads to better results.

**Table 2:** Comparison of predictions with different interaction analysis tools. The performance metrics used to compare PLIP, ProLIF, and po-sco are the PCC, the MUE, and the RMSE. The shown metrics apply to the external test set only. The best values are highlighted in bold font. All models were trained on a diverse split of the refined PDBbind set (see Section 3.1).

| Tool | PCC | MUE | RMSE |
| --- | --- | --- | --- |
| PLIP | 0.37 | 2.20 | 2.71 |
| ProLIF | 0.53 | 2.03 | 2.48 |
| Po-sco w/o L | 0.63 | 1.92 | 2.36 |
| Po-sco | **0.66** | **1.84** | **2.34** |

Both po-sco models with and without ligand information outperform the PLIP and ProLIF models, while the model with ligand information performed the best. This improved performance is likely due to the very high level of detail provided by posco. Not only does po-sco provide fine details about individual interaction types (e.g. hydrogen bond donor/acceptor in backbone/side chain), but it also extends binary fingerprints by continuous values, allowing the storage of a higher amount of information. The results in Table 2 show that this attention to detail is beneficial for the prediction of binding affinities using a deep learning model.

Adding information on the ligand “happiness”, which describes the level of saturation of various kinds of interaction partners by protein counterparts, seems to also benefit the model. However, it must be noted that in order to implement the ligand information, the prediction model used a slightly adapted architecture (see Section 3.3).

### 2.3 Impact of dataset split

The second goal of this work was to investigate the influence of dataset splitting, specifically the PDBbind dataset, on model performance. Here, we prepared three different splittings, two based on the complete PDBbind dataset and one based on the refined set. See Section 3.1 for details on the construction of the data set.

In a PDBbind splitting that is often seen in the literature, the general set is used for training, the refined set for validation, and the core set for testing. We analyzed the occurrence of proteins (i.e. Uniprot IDs) across the different sets in this split to identify potential overlaps. There were 2388, 1494, and 64 different proteins in the general, refined, and core set, respectively. Of the 1494 different proteins in the refined set, 74% were also part of the general set, and in the core set it was even 97%. Thus, even though there were different PDB entries in the training, validation, and test set in this split, there was a substantial overlap of proteins. This means that, while the exact conformation of the binding site residues likely differs in the different PDB entries of the same protein, the general similarity between different conformations of the same protein binding site introduces a large bias. This may allow a deep learning model to memorize the binding sites without learning much from the analyzed interactions and still perform well on similar binding sites.

To avoid such bias, we created a custom split that randomly assigns complexes to the training, validation, and test set, while avoiding spreading complexes containing the same protein across different sets.

We trained three versions of the same model. Model 1 was trained on the complete PDBbind dataset with the “classical” splitting, i.e. the general set for training, the refined set for validation, and the core set for testing. Model 2 was also trained on the complete PDBbind dataset, but with our custom split that enforces diverse proteins between sets. Model 3 was trained on the refined set only with the same diversity-optimized splitting as in model 2. Due to the best performance of the po-sco based prediction model compared to the ones based on PLIP and ProLIF, we focus here on models trained with po-sco inputs only. Data for the same analyses performed with PLIP and ProLIF-based models can be found in the Supporting Information in Tables S1 and S2, respectively.

Table 3 shows the performance of the model trained on the different datasets. It can be seen that, with respect to the prediction error, model 1 performs better on the test set than the other two models. However, this seemingly high performance is misleading, since 97% of the proteins in the test set were already seen during training. After removing overlapping proteins between training, validation, and test set, the performance drops significantly, indicating that the model has much more difficulty actually learning from the data (model 2).

**Table 3:**
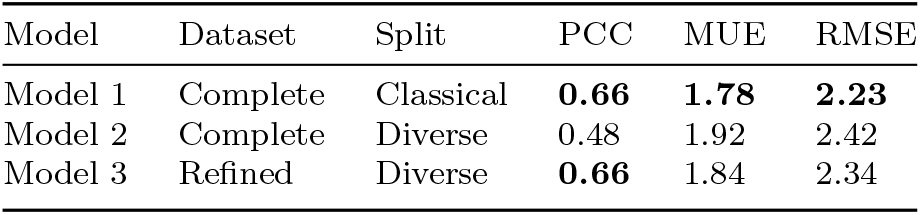
Comparison of the po-sco-based prediction model using different splittings of the PDBbind dataset. The performance is shown only for the external test set. The best performances are highlighted in bold.

Finally, when using only the refined set with a diverse split (model 3), the performance is higher compared to training on the complete dataset with the same splitting. Although the training set of model 2 was more than twice the size of that of model 3, the additional data points were not beneficial in model training. This indicates that for such tasks, the higher quality of the structure data in the refined set is clearly beneficial for meaningful (generally applicable and transferable) learning of the models.

### 2.4 Real-world example

We decided to test the trained models on a real-world example. For this, we chose the human 5HT receptor 1B (Uniprot ID P28222) because it is a pharmaceutically relevant target and is not part of the PDBbind dataset.

We docked more than 700 compounds with known binding affinities to the 5HT receptor using smina and Glide [12, 15]. The generated poses were re-scored with the three trained po-sco models and with gnina [5]. Again, we calculated the PCC, MUE, and RMSE of the different methods. The exact details can be found in Section 3.4.

Table 4 shows the results of this study. No tested method yielded satisfactory results, as the correlations between predicted and true affinities were quite weak. Judging by the MUE and RMSE, gnina does the best job in predicting binding affinities. Based on the performance reported in Table 3, it could be expected that model 1 performs the best out of all po-sco models. However, it in fact performed much worse than model 3 which was only trained on the refined subset of the PDBbind dataset. We think that model 1 mainly learned to memorize the structures in the training set. Due to the high overlap between the training, validation, and test set, the reported performance is accordingly high. When presented with a new protein structure, the model has difficulty in recognizing it and is unable to make an accurate prediction. Model 3, in contrast, was trained on data of superior quality with no overlap between the training, validation, and test sets, thus requiring it to gain more knowledge from the data itself. This clarifies why this model is more effective than model 1, even though the results in Table 3 may suggest otherwise.

**Table 4:**
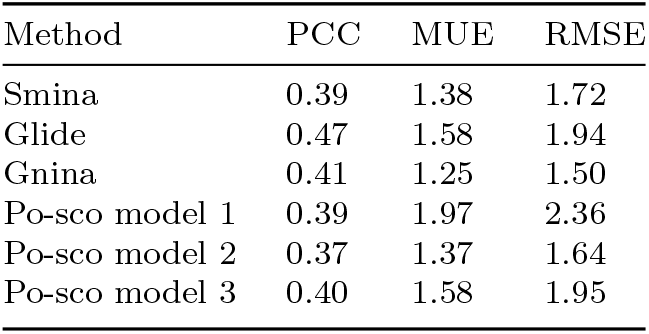
Comparison of different methods for the prediction of binding affinities towards the human 5HT 1B receptor (Uniprot ID P28222). The po-sco models 1, 2, and 3 were trained on the complete PDBbind dataset with a classical split, a diverse split, and on the refined subset with a diverse split, respectively.

Interestingly, model 2 which was trained on the complete PDBbind dataset with a split that removed all protein overlaps between training, validation, and test set, performed the best of the three po-sco models in this study. At first glance, this is surprising since the performance on the test set (Table 3) was much worse. One explanation could be that there are differences in the training or test set between models 2 and 3, leading to results that cannot be compared. However, 85% of the complexes that are in the training set of model 3 are also in the training set of model 2. For the test set, it is more than 75%. Also, as the protein at hand is not part of the PDBbind dataset at all, these effects will likely not have a big impact. Another possible explanation is that the performance on the test set as reported in Table 3 was a fluke and the true performance is better. This can be ruled out because the model performs equally as bad (or even worse) on the validation set (PCC 0.51, MUE 2.04, RMSE 2.47). A third explanation could be that the high performance of model 2 in this example was a “lucky shot” and is not representative for the general performance of the model which is better represented by the results in Table 3. Further investigation showed that while the human 5HT 1B is not part of the PDBbind dataset, the general set contains the turkey 5HT 1B receptor. In fact, this protein was part of the training set of models 1 and 2. Visual inspection showed that the binding sites of the turkey and human 5HT 1B receptors are not the same, although they are very similar. Thus, model 2 appeared to have an easier job predicting these compounds than model 3 because it has been trained on a very similar structure, which explains the higher performance. The fact that model 1 was also trained with this structure but still performed the worst of the three models further highlights that it probably did not have to learn actually meaningful information to still achieve a seemingly high performance due to the overlap between the data sets.

Another point that needs to be kept in mind is that in this experiment, the model was applied to poses from molecular docking whereas it was trained on crystal structures. This could also have an influence on the outcome.

We strongly advocate that researchers be very mindful about the data set used to train a model and how it was handled. This enables the early detection of reported performances that have been exaggerated and prevents the scientific literature from being inundated with models that look good on paper, but fail in real-world scenarios. It is essential to critically evaluate the reported performance of a model.

## 3 Methods

### 3.1 Dataset construction

As described in Section 1.4, the protein and ligand structures provided in the PDBbind dataset need to be prepared before using them for structure-based modeling approaches. We used the protein preparation wizard in Schrödinger’s Maestro to prepare all structures [64]. The preparation workflow consisted of adding all explicit hydrogen atoms, assigning bond orders, generating protonation states at the pH present at crystallization (for the protonation of the ligand, Epik was used [65–67]), generating water orientations, and running a restrained minimization with an RMSD cutoff of 0.3 Å.

In this work, we used several different splits of the PDBbind dataset version 2020 (Figure 2). The complete dataset (consisting of the general, refined, and core set) contained 19,443 protein-ligand complexes. For the general set, there were affinities of Ki, Kd, and IC50 types. Since Ki/Kd and IC50 cannot be directly compared, we removed samples with affinities of type IC50. This left us with 6558 complexes in the general set, 5041 in the refined set, and 256 in the test set after removing all duplicate complexes.

**Fig. 2:**
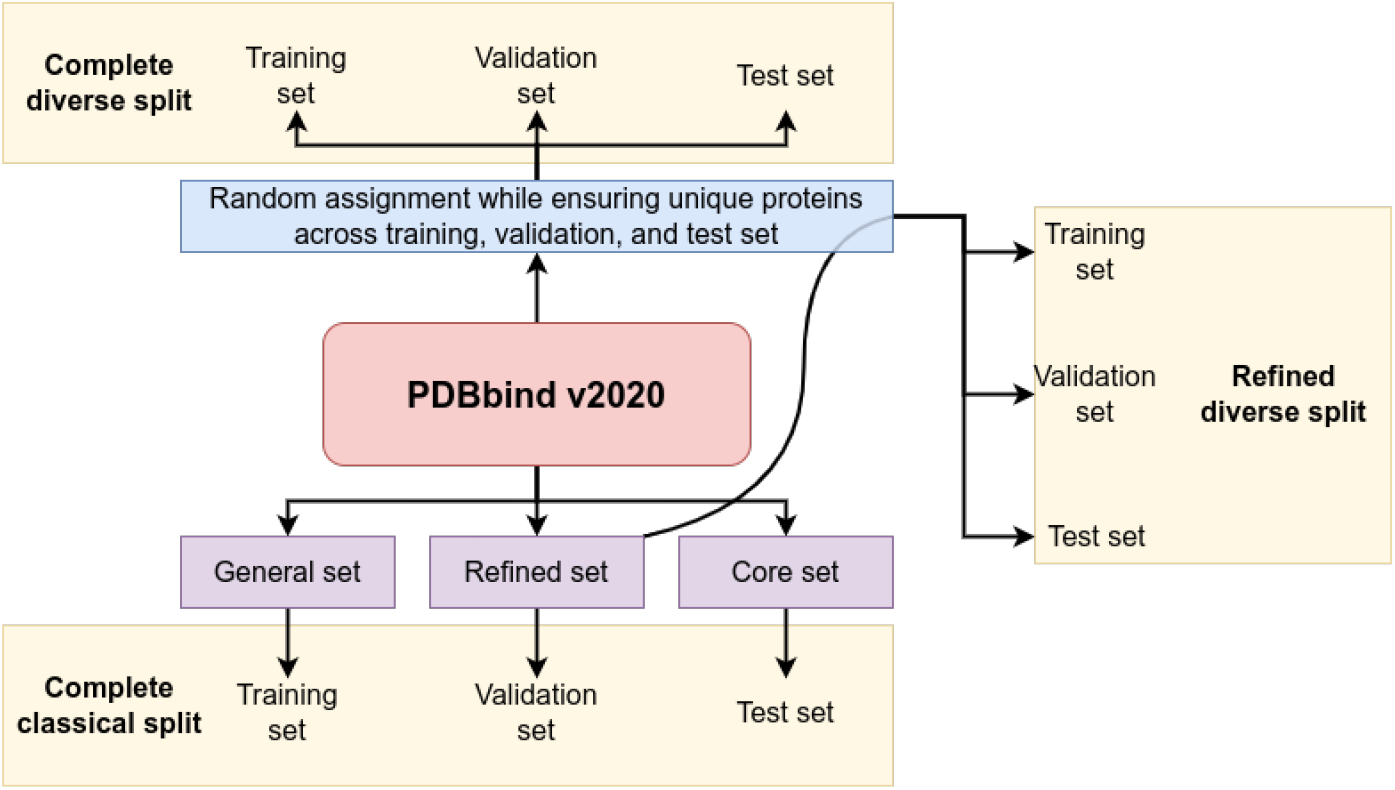
The splitting scenarios applied to the PDBbind dataset. The complete dataset was split in two ways, one in which the general, refined, and core set are used for training, validation, and testing, respectively (complete classical split), and one in which the complexes were split randomly while ensuring that there are no overlaps of proteins between the training, validation, and test set (complete diverse splitting). The same diverse splitting was applied to the refined set only (refined diverse splitting).

The complete dataset was split in two ways: 1) The classical split, where the general set was used for training, the refined set for validation, and the core set for testing; 2) the diverse split, where the complexes were randomly assigned to either the training, validation, or test set while ensuring that there are no overlaps regarding the protein (i.e. Uniprot ID) between the datasets. This means that, while there were several complexes containing the same protein in one dataset, the proteins were unique between different datasets. The training set contained around 80% of the complexes, while the validation and test set contained around 10% of the complexes each. For the refined set (containing complexes of higher quality compared to the general set), the same technique for generating a diverse split was used.

The model architecture used here, combined with the provided input, does not allow conclusions to be drawn about the structure of the ligand. This was specifically chosen to avoid the introduction of bias and thus prevent the model from simply memorizing structures of ligands.

### 3.2 Po-sco interaction analysis

The po-sco interaction analyzer tool contains algorithms to detect the most significant intermolecular protein-ligand interactions currently recognized by the modeling community. The list of all supported interaction types can be found in Table 1.

Detection algorithms have been implemented in such a way that they reliably identify specified interactions within reasonable structural deviations (see below). While we trained our models on crystallographic data, where realistic interaction patterns (e.g. interatomic distances, angles, ring and π-system planes) are naturally expected, we aim at using po-sco for scoring of docked poses, where moderate deviations from ideal parameters can be frequently observed, even with “good” poses that we typically refer to as those within 2.0 Å from the native pose. Therefore, e.g. for distance thresholds, a general factor allowing for a 20% deviation from the optimum is applied. In terms of linearity, a deviation of up to 45 degrees is accepted.

Compared to similar tools, in many cases, our routines are parameterized with more stringent criteria. For example, for detecting pi-pi face-to-face stacking and T-shape interactions between rings we use different cut-off values for centroid distances, which helps minimizing false positive as well as false negative cases. We implemented our expertise from modeling and docking small molecules to a multitude of different targets covering membrane bound and soluble proteases, metalloenzymes, cytochromes, GPCRs, nuclear receptors, ion channels, etc. During the development we relied heavily on benchmarking with real-world systems. The H-bonding parameters, both in terms of geometry and energetics, are based on the Yeti force field by Vedani et al. [68]. The same applies for ligand-metal interactions [69].

The algorithm for the atom-type-dependent quantification of hydrophobic interaction was parameterized based on the relative solvation free energies of matched molecular pairs of unsubstituted benzene, toluene, fluro-, chloro-, bromo-, iodobenzene and thiophenol in the organic solvent hexadecane (cf. Figure S3). The underlying data were extracted from the Minnesota Solvation database [70]. Toluene was selected as the reference substance, providing a weighting coefficient of 1.0 for the sp3-hybridized carbon atom. Equation 1 shows the calculation of the hydrophobic interaction energy between atoms *a_i_* and *a_j_* where *w_i_* and *w_j_* are the weights according to Table S3 for the respective atoms, *k* is an additional factor according to Li et al. [71], *r_vdw_*(*·*) is the Van der Waals radius of a given atom, *n* is a slope parameter, and *d_ij_* is the distance between atoms *i* and *j*. The slope parameter *n* was set to *−*6.0 and following the article by Li et al., we defined *k* = *−*0.3.

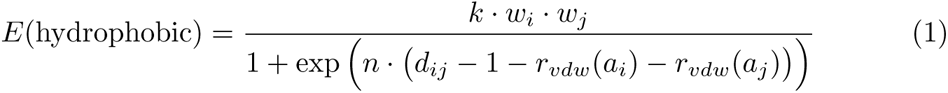

Ring stacking interaction is detected up to the ring centroid distance of 5.0 Å. Depending on the ring character (aromatic or aliphatic), a detailed analysis of ring interaction patterns is performed. In case of aromatic rings, the pair-wise distances are calculated between all atoms in the first ring and all atoms in the other ring counting the number of hydrogen and heavy atoms at or below the sum of the Van der Waals radii. Based on the prevailing interactions (hydrogen-to-heavy or heavy-to-heavy), the type of interaction is determined (T-shaped or parallel). In our experience, this procedure works more reliably than calculating the angle of ring planes.

Detection of complex interaction patterns, e.g. polarized H atoms interacting with an aromatic ring, is based on a thorough analysis of the local environment of interacting atoms, i.e. for switching the relevant fingerprint bit on, such a polar hydrogen atom must be within acceptable interaction distance (below the sum of VdW radii) with at least three heavy atoms forming the target pi system of the ring, and simultaneously the hydrogen exit vector must be co-linear (with the ring normal with a maximum allowed deviation, here 30 degrees).

Similarly, pi-pi stacking of smaller, non-ring systems is detected for at least three consecutive sp2 hybridized atoms (e.g. peptide bond π-system). After confirming the validity of geometric criteria, interactions are classified according to the chemical character of atoms in higher order interacting functional groups (neutral, positively or negatively charged π-system and all combinations thereof).

Halogen bonding is also detected at interatomic distances below or at the sum of Van der Waals radii of the analyzed atom pair. Depending on the angle with respect to the atom, which carries the halogen atom, interactions are classified to the sigma hole (angle carrier atom - halogen - heteroatom is greater than 135 degrees) or the belt (angle carrier atom - halogen - hydrogen in the range of 45 to 135 degrees). Halogen sigma hole - aromatic ring interactions are detected if at least two ring atoms engage with the halogen atom and the ring normal is within 45 degrees of the halogen sigma hole bond vector.

Some geometric parameters (e.g. H-bond donor or acceptor saturation, and happiness of hydrophobic atoms) can be correctly evaluated only by considering their buriedness, i.e. accessibility for interaction with the bulk water. Buriedness is calculated for each ligand and protein atom. First, a grid of points (1.0 Å spacing) is generated around the ligand extending 12 Å ligand extremes in each Cartesian axis direction (-x, +x, -y, +y, -z/+z). Next, all grid points within the distance below 3.0 Å from either protein or ligand atoms are deleted. Finally, the remaining points are analyzed for their vicinity to ligand and protein. Atoms appearing within the threshold value of 6.0 Å from any grid point are marked as not buried.

### 3.3 Attention-based prediction model

The po-sco prediction model uses three outputs from the po-sco interaction analyzer as input. The first output is the type of interacting residues, the second output is the interaction fingerprint between protein residues (including waters and metals) and the ligand, and the third output is the ligand “happiness” fingerprint describing the level of saturation of interaction partners in the ligand molecule by the protein counterparts. Here, we describe the architecture of the prediction model based on po-sco (Figure 3).

**Fig. 3:**
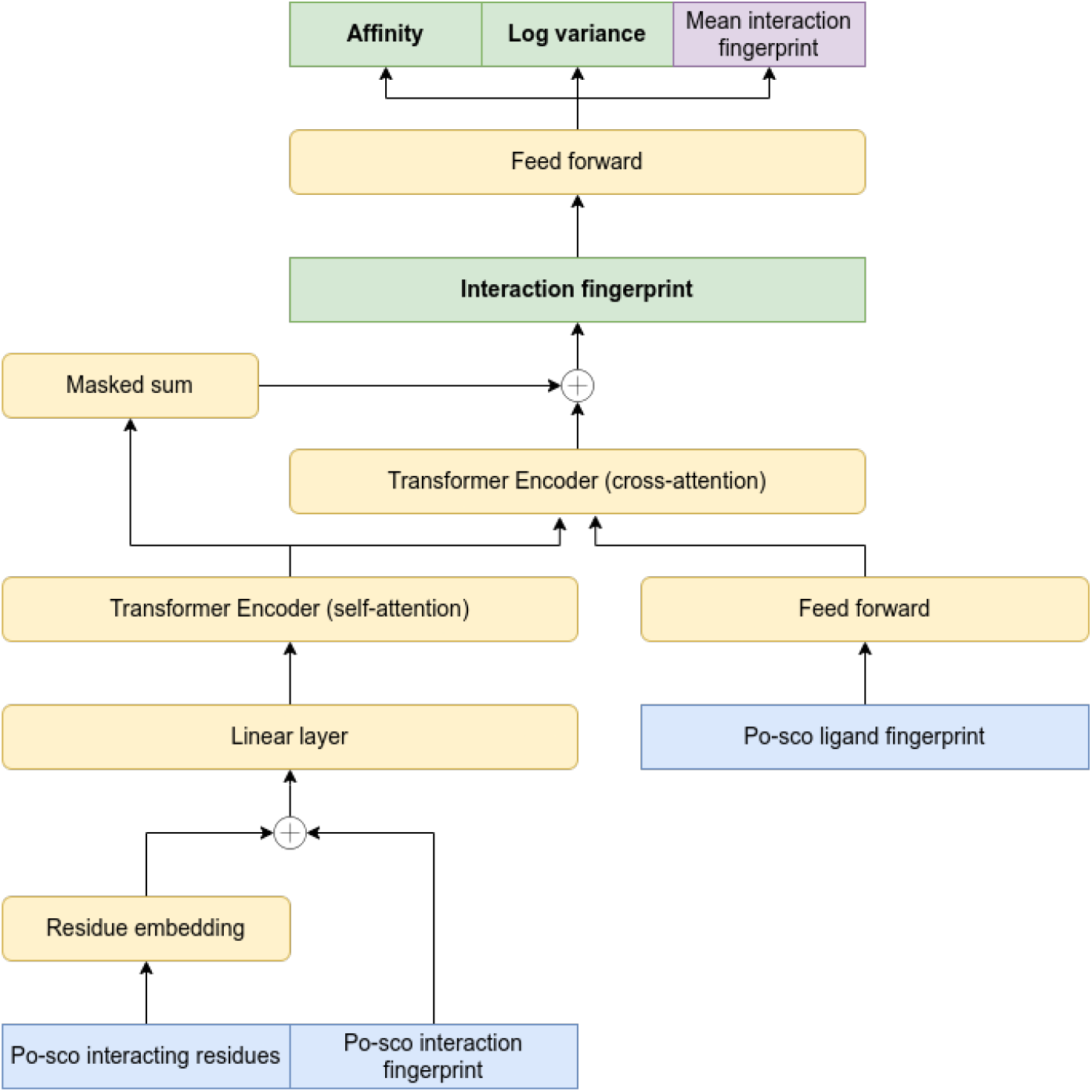
The architecture of the prediction model. Blue: Output obtained from the posco interaction analyzer. Yellow: Neural network layers. Green: Usable model outputs. Purple: Only used for training to increase the conservation of interaction information.

The type of interacting residues is first embedded in an 8-dimensional space and concatenated with the corresponding interaction fingerprint. This combined input is then passed through a linear layer to embed the data in a 64-dimensional space. The embedded data are then used to compute self-attention with a transformer encoder layer.

The ligand fingerprint is also embedded into a 64-dimensional space using a fully connected layer. The embedded ligand is then used to calculate cross-attention with the encoded residue information using a transformer encoder layer.

To conserve information on the interactions in the complex, we added the encoded residue information to the output from protein-ligand cross-attention. Since the output from the cross-attention layer is a vector, whereas the encoded residue information is a matrix, we applied a masked sum on the encoded residue information over the first dimension of the matrix. The mask is used to mask out padding elements in samples of unequal length. The output of this sum operation can be used as a non-binary single vector interaction fingerprint describing the protein-ligand complex. This vector could also be used for down-stream predictions other than the binding affinity.

The generated interaction fingerprint is then passed through a feed forward network to predict the affinity, the log variance of the affinity, and the mean po-sco interaction fingerprint. The predicted affinity and log variance are used to train the model using maximum likelihood estimation. In addition to the maximum likelihood estimation, the mean po-sco interaction fingerprint can be used to train the reconstruction of the interactions, which is intended to increase the amount of information (about the interactions in the complex) stored in the predicted interaction fingerprint. This reconstruction is trained using a mean squared error loss. The complete loss function is therefore a combination of a negative log likelihood loss and a mean squared error loss:

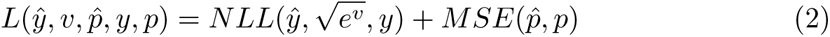

Where 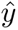 is the predicted affinity, *y* is the true affinity, *v* is the predicted log variance of the affinity, 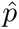 is the predicted mean po-sco interaction fingerprint, and *p* is the true mean po-sco interaction fingerprint. The negative log likelihood loss is defined as:

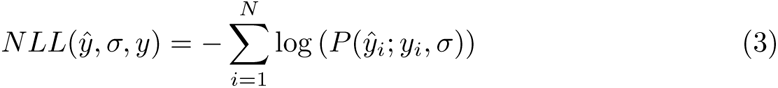

where 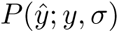 is the probability density function, i.e. the probability of finding 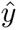 under a normal distribution given by the mean *y* and standard deviation *σ*, defined as:

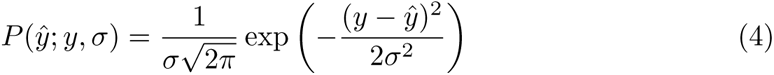

The mean squared error loss is defined as:

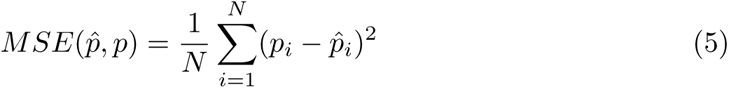

The model was trained in PyTorch using an Adam optimizer [72]. The model was trained with 2 transformer encoder layers for the self- and cross-attention each, an embedding dimension of 64, 2 heads per attention layer, a batch size of 64, a learning rate of 1*e^−^*^4^, and a dropout rate of 0.15. All models were trained for 1000 epochs, and the epoch with the lowest validation loss was used to assess the model’s performance (on the test set). All models were trained with the same hyperparameters where applicable.

#### 3.3.1 Ablation study

In order to test the contribution of the ligand fingerprint calculated by posco, we performed an ablation study in which the ligand fingerprint was not used in model training. In this model, the predicted interaction fingerprint was calculated directly as the masked sum of the encoded residue information. Thus, there was no cross-attention with the ligand. The rest of the model remained the same.

#### 3.3.2 Model based on PLIP

Since PLIP is commonly used to calculate protein-ligand interactions, we trained a model using the information obtained from PLIP. In our tests, we used PLIP version 2.2.2. The interactions calculated using PLIP can be found in Table 1.

The architecture of the model used with PLIP followed that of the one used in the ablation study, because no ligand fingerprint was used.

#### 3.3.3 Model based on ProLIF

We trained an additional model using ProLIF version 1.1.0, one of the other tools commonly used for the analysis of protein-ligand interactions. The interaction types used from ProLIF can be found in Table 1. Like the PLIP-based model, the architecture of this model followed that in the ablation study without ligand information.

### 3.4 Real-world example

We downloaded compounds with known activity toward the human 5HT 1B receptor (Uniprot ID P28222) from PubChem [73]. The obtained substances were further filtered to exclude all samples that do not have an activity type of Ki or Kd, that do not have an exact affinity value, or that do not have unambiguous stereochemistry. This left us with a total of 726 compounds. Before docking, all compounds were prepared with LigPrep to generate protonation states at pH 7.4 [74].

The compounds were docked with smina against the PDB IDs 4IAQ, 5V54, and 6G79 and with Glide against the PDB IDs 5V54, 6G79, and 7C61. For each compound, up to 9 poses were generated. For Glide, the Glide SP protocol was used, and the binding site was defined based on the location of the co-crystallized ligand. For smina, the binding site was defined based on the co-crystallized ligands with a default buffer of 4 ^Å^ on all six sides and the compounds were docked with an exhaustiveness of 16 and a seed of 42.

All protein structures were prepared by adding explicit hydrogen atoms, assigning bond orders, converting selenomethionines to methionines, filling in missing side chains and loops, generating protonation states at pH 7.4, optimizing the water network, and performing restrained minimization with an RMSD cutoff at 0.3 Å. All preparation was carried out in Schrödinger’s Maestro using the protein preparation wizard [64].

The generated docking poses were re-scored with gnina and three trained po-sco models. Gnina was used with the --score only flag, 2 CNN rotations, a seed of 42, and an exhaustiveness of 16. The best predicted score for all poses of a compound was used to calculate the performance statistics.

## 4 Conclusion

Recently, large language models trained on large amounts of data have gained increasing attention [39, 75, 76]. In this work, we show that the quality of the data used to train a model is more important than the quantity when it comes to the prediction of binding affinities based on molecular interactions. We demonstrate how overlaps between the training, validation, and test set, specifically in the PDBbind dataset, can lead to severe overestimation of model performance. We also suspect that many recently published models for binding affinity prediction may suffer from such a bias. In this regard, we urge researchers to critically assess the data used to train deep learning models and to avoid bias introduced by poor dataset splits. We also suggest working with smaller datasets of very high quality rather than large datasets of poor quality.

Furthermore, we introduced po-sco, a sophisticated tool for the analysis of protein-ligand complexes that provides a great level of detail. We showed how the high content of information extracted by po-sco benefits deep learning models. A transformer-based model that utilized analyses from po-sco was more successful in predicting binding affinities than models that were trained with data from PLIP and ProLIF. We propose that a very detailed and physics-based analysis of protein-ligand interactions allows deep learning models to better learn from structural data and improves downstream predictions.

Finally, we promote the use of structure-agnostic models, i.e. models that do not explicitly know the structure of the ligand or binding site. In this way, the risk of creating biased models can be reduced to a minimum.

## Acknowledgments

We gratefully acknowledge the support of NVIDIA Corporation with the donation of two RTX A5000 GPUs used for this research.

## Notes

### Competing Interest Statement

The authors have declared no competing interest.

