## Supplementary Information for "Quality Matters: Deep Learning-Based Analysis of Protein-Ligand Interactions with Focus on Avoiding Bias"

**Table S1**: Comparison of the PLIP-based prediction model using different splittings of the PDBbind dataset. The performance is shown only for the external test set. The best performances are highlighted in bold.

| Model | Dataset | Split | PCC | MUE | RMSE |
|---|---|---|---|---|---|
| Model 1 | Complete | Classical | **0.48** | **2.05** | **2.56** |
| Model 2 | Complete | Diverse | 0.18 | 2.14 | 2.68 |
| Model 3 | Refined | Diverse | 0.37 | 2.20 | 2.71 |

**Table S2**: Comparison of the ProLIF-based prediction model using different splittings of the PDBbind dataset. The performance is shown only for the external test set. The best performances are highlighted in bold.

| Model | Dataset | Split | PCC | MUE | RMSE |
|---|---|---|---|---|---|
| Model 1 | Complete | Classical | **0.57** | **1.91** | **2.41** |
| Model 2 | Complete | Diverse | -0.03 | 2.16 | 2.71 |
| Model 3 | Refined | Diverse | 0.53 | 2.03 | 2.48 |

**Table S3**: Weight factors for the calculation of hydrophobic contributions. The weight $w$ of a solute $x$ is calculated as $w = \frac{\Delta G_s(x)}{\Delta G_s(\text{Me-benzene})}$ where $\Delta G_s$ is the free energy of solvation of solute $x$ in the organic solvent hexadecane and Me-benzene is toluene.

| Solute | $\Delta G_s(hexadecane)$ | weight |
|---|---|---|
| Benzene | -3.80 | 0.837 |
| Me-benzene | -4.54 | 1.000 |
| F-benzene | -4.03 | 0.888 |
| Cl-benzene | -4.99 | 1.099 |
| Br-benzene | -5.51 | 1.214 |
| I-benzene | -6.25 | 1.377 |
| SH-benzene | -5.61 | 1.236 |

**Acknowledgments.** We gratefully acknowledge the support of NVIDIA Corporation with the donation of two RTX A5000 GPUs used for this research.
